# Modelling the effects of land use on mangroves in a RAMSAR site of Panamá

**DOI:** 10.1101/2021.10.07.463581

**Authors:** Juliana López-Angarita, Juan M Diaz, Alexander Tilley

## Abstract

The role of mangroves as pivotal providers of ecosystem services has been widely acknowledged. In Latin America, mangroves play an important role in traditional coastal livelihoods, but the growing economy of these nations demands the expansion of land for development, putting pressure in ecosystems such as mangroves. Here we examine the impact of land use activities on mangroves in the Gulf of Montijo, a RAMSAR site located in the Pacific coast of Panamá. Spatial information of land use was analysed, ground-truthed and classified into agriculture, aquaculture and coastal development, and subsequently ranked according to estimated level of impact on mangroves based on 27 interviews with local informants. We developed a spatially-referenced cumulative impact model of human activities on mangroves. Results showed that despite the protection status of the Gulf of Montijo, its mangrove forests are affected by localised human activities, dominated by agriculture. Given the importance of fishing for local livelihoods, evaluating the effects of agriculture, rice in particular, on mangroves and their associated fauna will be essential for the sustainable management of this RAMSAR site.

## 1. Introduction

In Central America, mangroves were utilized by pre-Columbian people around 5000 years ago for timber, charcoal, tannins, and fishing, with Ariidae, Carangidae, Clupeidae, Sciaenidae, Batrachoididae particularly targeted (Cooke and Ranere 1999). Mangroves are of direct livelihood importance to fishing communities living along Panamá’s Pacific coast (Trejos et al. 2007a; Trejos et al. 2008). However, Panamá is one of the countries that has experienced the highest mangrove deforestation rates in the eastern tropical Pacific region, with around half lost in the last 50 years (FAO 2007; ANAMARAP 2013; López Angarita et al. 2016). Agriculture, livestock, salt production, aquaculture, urban or industrial development, and direct extraction for tannins and wood, have been identified as main drivers of deforestation (D’Croz 1993; Ibáñez et al. 2005).

Although Panamá ratified the RAMSAR Convention of Wetlands in 1989, it wasn’t until a decade later that the country’s first mangrove protection appeared in legislation (Cooke and Ranere 1999; López Angarita et al. 2016). In 2008, all Panamá’s mangroves became marine-coastal management areas, where logging, and any use or commercialization of the system is prohibited, except in areas subject to special regimes, such as sustainable resource management zones (ANAMARAP 2013). The Panamanian government allows commercialization of mangrove products (e.g. charcoal, timber, tannins) in permitted areas via harvesting permits (e.g. the Gulf of Chiriquí), however, the system is poorly enforced and unsustainable harvesting occurs (Trejos et al. 2008). While pond construction for aquaculture is prohibited in all mangrove areas, it is allowed in tide flats or “albinas” with a special permit, and once a shrimp farm is established, ‘minor alterations’ can then be made to the surrounding forests to install water pumps, channels and sluice gates (Maté 2005). Similarly, mangrove clearance is allowed for tourism development (Law 2. Jan 7, 2006). Such measures clearly contradict the idea that marine resources are national patrimony in Panamá (Law 44, Nov 23, 2006) (Castellanos-Galindo et al. 2017).

The Gulf of Montijo (GoM), located in the south of Veraguas Province on the Pacific coast of Panamá (Figure 1), is one of the most important mangrove systems of the country and was designated as a RAMSAR site in 1990, spanning 80,765 ha (RAMSAR 1990). The GoM was declared a protected area in 1994 and holds the management category of “Wetland of International Importance” (Pinto and Yee 2011). Local communities are primarily engaged in fishing, agriculture, and tourism (Pinto and Yee 2011; Ventocilla 2013). Last population census in 2010 estimated the population of the GoM to be around 6572 inhabitants (INEC 2011). The Montijo wetland is complex with many river deltas, beaches, rocky shores, mudflats and mangroves. The wetland is an important habitat for migratory aquatic birds and large mammals such as monkeys and sloths (CREHO 2010), but it is also a very important agricultural region with a large amount of fields dedicated to crop production, mainly rice, in its flood plain (ANAM 2004). Artisanal fisheries of the GoM target finfish, sharks, lobster and shrimp, using mainly gillnets (Vega et al. 2014). In 2012, the government registered 25 artisanal fisheries cooperatives, with 499 associated fishers and 214 boats operating in the GoM (Vega et al. 2014). For the same year, Panamá’s fisheries department confirmed there were 776 active fishing licences for the GoM (Vega et al. 2014).

**Figure 1.**
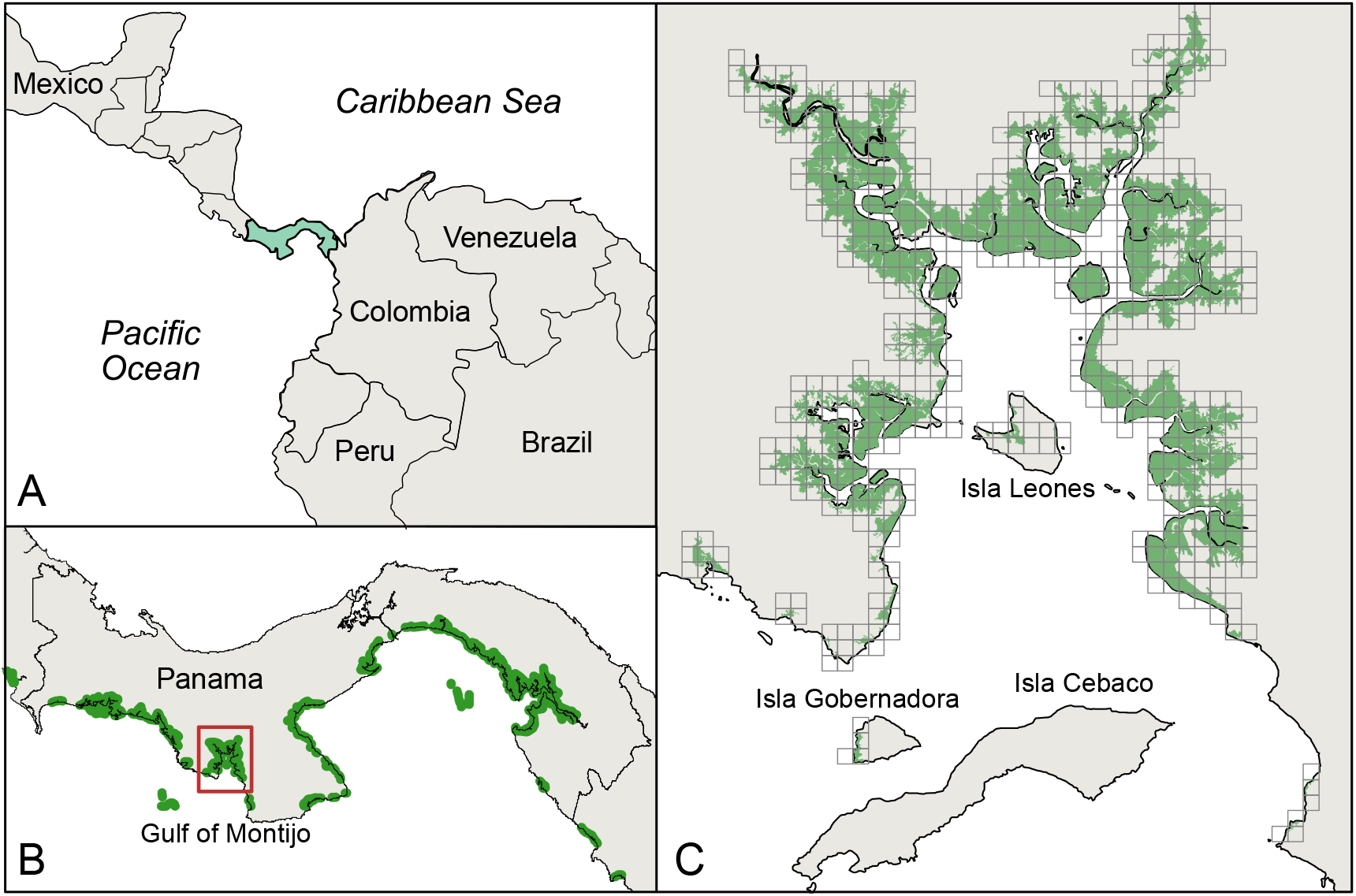
(A) The location of Panamá in Central America, and (B) the location of the Gulf of Montijo in Panamá (inside square). (C) The Gulf of Montijo divided in 1 km^2^ sample units used to classify impacts along the Gulf’s coast. Green colour in B and C represents mangrove forest.

The management of dynamic wetland landscapes such as the GoM, where multisector interests are the common, requires understanding the social and ecological drivers of change such as the spatial patterns of land use and land cover change, or the degree of dependence on mangrove resources by local communities. In this study, we explore land use adjacent to mangroves in the GoM using satellite images and ground-truthing, and develop a cumulative model of anthropogenic impacts as a step towards understanding the complexities of this wetland. We aim to provide managers with information essential for spatial planning in an exercise for supporting management for this RAMSAR site.

## 2. Methods

### 2.1. Interviews

We visited 12 coastal villages in the GoM where fishing was a common livelihood and performed interviews with key informants about mangrove condition. We decided to interview people related to fishing as this activity closely associated with mangroves and people interacted with mangroves day to day. To guarantee respondents were knowledgeable about mangroves, we interviewed experienced fishers (i.e. older than 50 years of age). Interviews were designed with open questions to gauge the community perception of mangrove condition and their contribution to local fisheries (Supporting Information). Interviews were only carried out after obtaining the verbal consent of each person. All our activities in the GoM were carried out under the guidance of Marviva Foundation, who has extensive experience working with fishing communities in the area, and under the respective field research permits issued by the Panamá Government Ministry of Environment. Questions focused on the respondents’ perception of: mangrove health and changes in coverage over time; the identification of human impacts affecting mangroves; the role of mangroves supporting fisheries; and the importance of mangroves for their livelihoods. Interviews were recorded with respondent’s permission and transcribed to perform qualitative analysis using the software Nvivo 10.2. Results were analysed by calculating the proportion of respondents with similar answers per question according to coded topics.

### 2.2. Identifying human activities adjacent to mangroves and impact model

We identified and classified human activities and impacts around the GoM using spatial information of land use obtained from the Marviva Foundation, the Panamá Government Ministry of Environment and the Smithsonian Tropical Research Institute (Supporting information 1). Given the information available, we grouped human activities into three major classes: aquaculture, agriculture, and populated centres. Land use classification was verified using Google Earth and through ground-truthing. We ground-truthed locations with repetitive features in satellite images, as well as features that were blurry or of low resolution. Once in the field, we went to the identified places in person, or if they occurred on private property or were inaccessible we visually verified and estimated the ground cover and any impact from a nearby vantage point, or contacted the land owner to verify land use. GPS points and *in situ* photographs were taken.

Human activities adjacent to mangrove forests pose direct or indirect threats to the ecosystem. To assess human impacts to the GoM’s mangroves, we built a model of cumulative impact following Halpern et al. (2008), where human activities that impact mangroves directly or indirectly were overlaid onto a map of mangrove coverage together with the ground-truthed maps of land use. The cumulative impact of these activities in mangrove forests was calculated in 1km^2^ sample units obtained by superimposing a 1km^2^ grid where mangroves were present in the GoM. Grid cell values were checked against the human activities identified during ground-truthing, correcting values where appropriate. Each 1km^2^ was given a score based on the presence (1) or absence (0) of each human activity in the cell. The model calculates a measure of impact by arithmetically adding the scores of human activities within each cell of the grid. We calculated a standard impact score by assigning each activity the same weighting, and a weighted impact score. The un-weighted cumulative impact score (CIS) was calculated for each 1 km^2^ cell as follows:

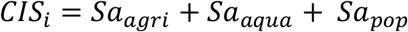

where *Sa* is the score of each human activity (1=presence and 0=absence), and ***i*** corresponds to each cell of the grid. Subscripts represent the first letters of each activity: agriculture, aquaculture, and populated centres).

For the weighted model, weightings were calculated using a ranking method adapted from Halpern et al. (2007), using scores of mangrove vulnerability to each human impact based on interviews with GoM fishers. Fishers were asked to identify the most important impact to mangroves of the area. Vulnerability scores were obtained for each impact using the frequency with which that impact was mentioned in responses to determine their relative importance. Once the frequencies for each human activity were identified, we ranked human impacts from highest to lowest impact (using numbers from 4 to 2). We use ranking instead of frequencies because we did not want to extrapolate trends from the different frequencies in our sample to all the GoM. In each 1 km^2^ cell, vulnerability scores for each human activity were summed according to their presence or absence. The weighted cumulative impact score (WCIS) was calculated for each 1 km^2^ cell as follows:

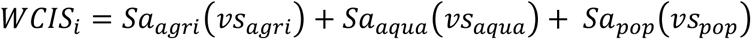

where *Sa* is the score of each human activity (1 = presence and 0 = absence); *vs* is the vulnerability score of the activity; and ***i*** corresponds to each cell of the grid.

For visualization purposes, the model attributes a colour to each 1km^2^ grid cell based upon the cumulative impact score, on a green to red scale, representing the state of mangroves according to the influence of adjacent human impacts. Finally, to quantitatively analyse the effect of land use on mangroves, we calculated the percentage of cells under each category of impact and under each category of human impact for both models.

## 3. Results

### 3.1. Interviews

Twenty-six men and one woman were interviewed from twelve fishing villages in the GoM (Supporting information). All were fishers who had lived their entire lives in the area. Eighty-nine percent were older than 40 and most had been fishing since they were young. 96% of respondents reported that fish productivity had declined in the GoM compared to when they started fishing, and 30% mentioned the collapse of snapper and shrimp fisheries specifically. 46% of respondents felt that overfishing was the main cause of fishery declines with natural fluctuations of fish abundance and climate change also mentioned by 22% and 7%, respectively.

Agriculture was the most important threat to mangroves raised by all respondents (100%), as this had caused their clearance in the past. Likewise, all stated that fumigation and fertilization of agricultural fields, during the dry season, and for rice in particular, caused massive die-offs of fish during the rainy season when rivers wash chemicals into the sea. When asked if they felt mangroves were important to their fishing, 100% of respondents said they were, and most (65%) were aware that this was due to their provision of food and habitat to juvenile fish (i.e. as nursery grounds). Snook (*Centropomus medius*), snappers (Lutjanidae), hammerhead sharks (Sphyrnidae), mullet (Mugilidae), weakfish (Sciaenidae), shrimp (Penaeidae), catfish (Ariidae), cockles (*Anadara tuberculosa* and *A. similis*), and crabs (*Callinectes arcuatus*) were identified as important commercial species that only occur around mangroves. 18% said mangroves help maintain water quality, and 15% said they provide aesthetic value to the landscape. Illustrative comments on these topics included:

> *“Mangroves are everything, as they give life to animals and humans.”*
>
> *“No mangroves, no fish*.*”*

### 3.2. Identifying human activities adjacent to mangroves and impact model

The main economic activities in the GoM are cattle farming for milk and beef, and rice production predominantly for the national market. Cattle graze in fields with plenty of native vegetation. In addition, on smallholdings, beans, sugarcane and banana are grown for local consumption, but the detail of these is not apparent on satellite images given their small size (*pers. obs*. 2014). The results of ground-truthing were used to verify the source maps of land use in 145 cells of 1km^2^ from a total of 632 cells from the grid placed over the GoM. A mangrove state model was generated to incorporate this information. Figure 2 shows the resulting overlay from the un-weighted impact score, with four levels of impacts ranging from very high (i.e. cells with all human activities present) to low (i.e. cells with only mangrove present and no human activities). For the weighted model, agriculture was the activity with the highest vulnerability score as all interviewed fishers mentioned it as the activity that impacted mangroves the most. Aquaculture was classified as the second most threatening activity and centres of population had the lowest vulnerability score (Table 3). Impact score in each cell was calculated according to integer values of the vulnerability scores. Categories of impact increased from four in the unweighted model, to seven in the weighted model, ranging from low to very high impact (Supporting Information).

**Figure 2.**
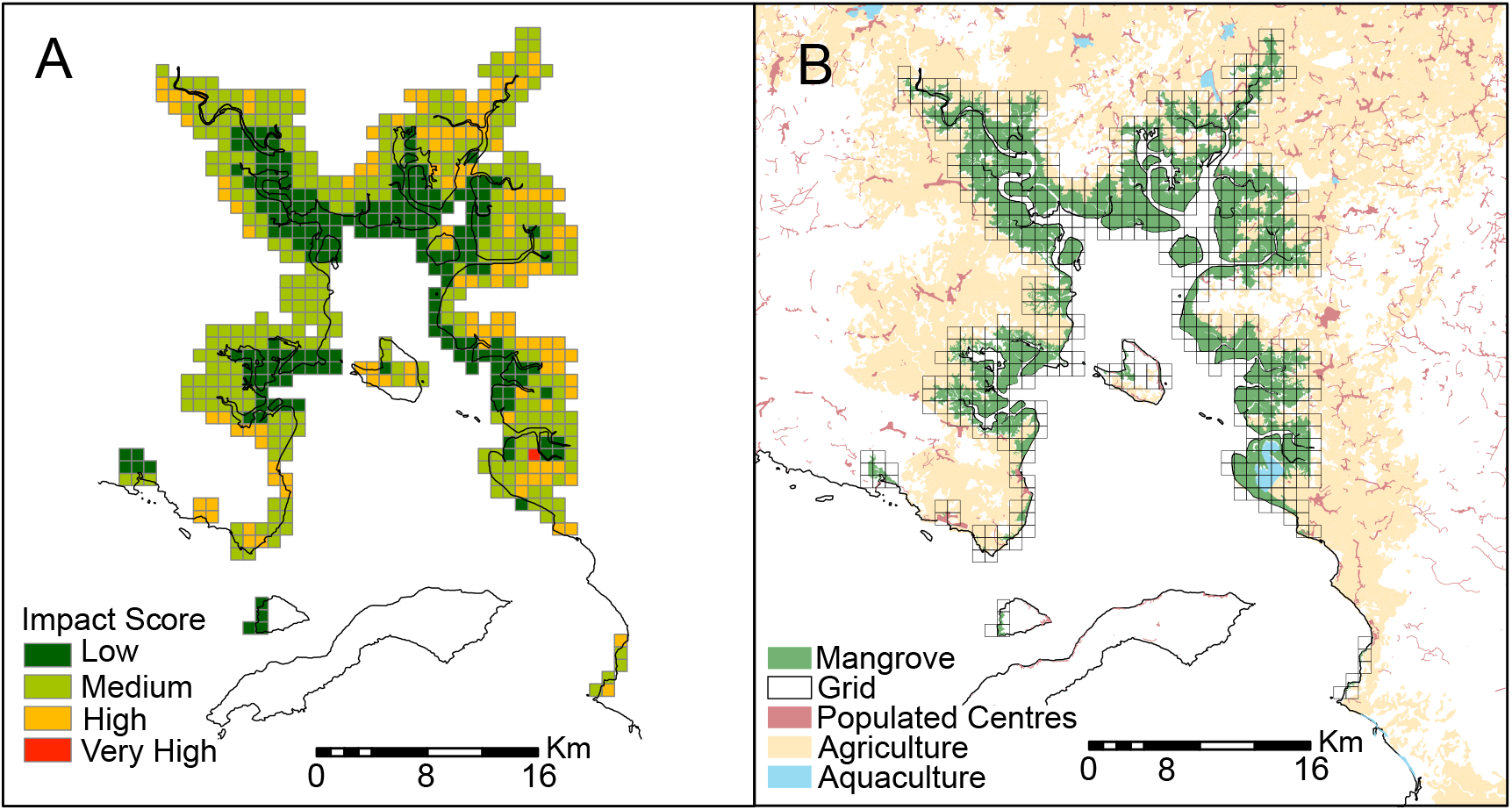
The Gulf of Montijo, Panamá. (A) Standard cumulative impact model for human activities per 1km^2^ in the Gulf of Montijo, showing mangroves with low impact in green and high impact in red. (B) The main human activities adjacent to mangroves are agriculture, aquaculture and centres of population.

**Table 1.**
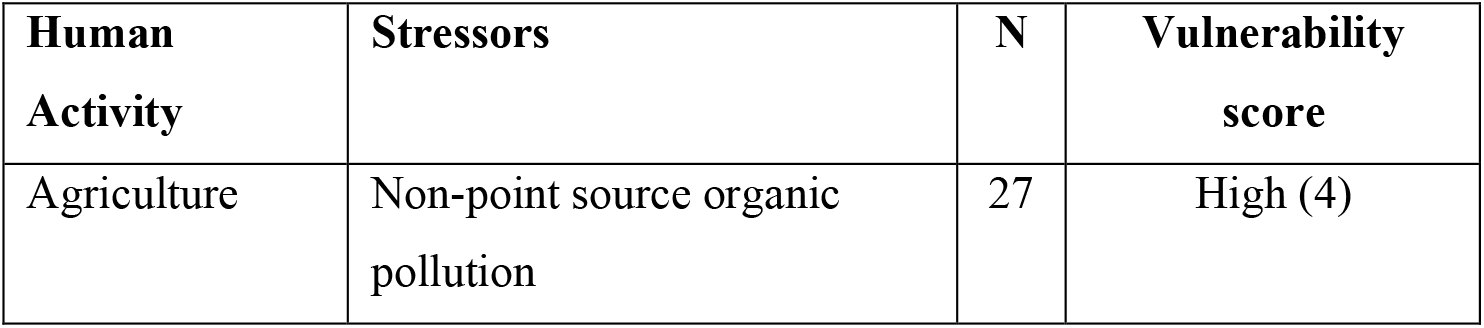
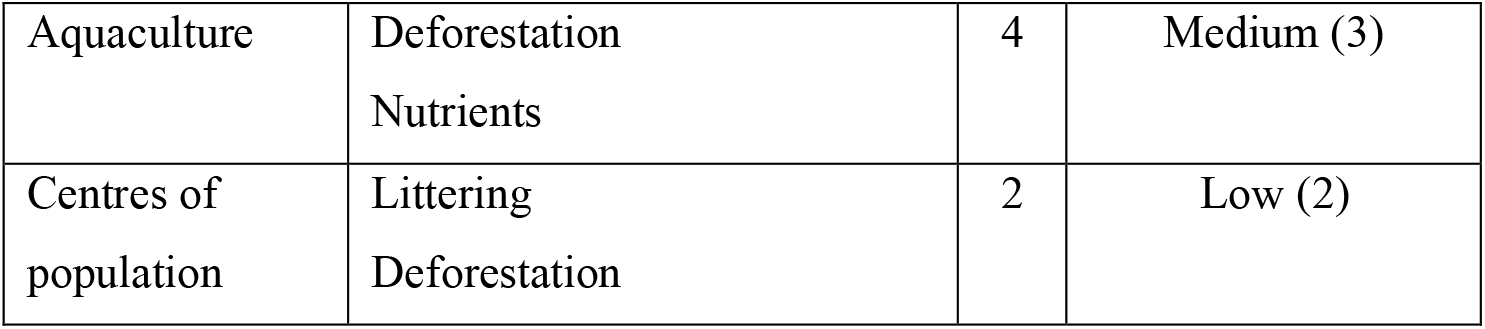
Scores given by Gulf of Montijo (Panamá) fishers to reflect mangrove vulnerability to threats from human activities. N refers to the number of respondents that mentioned a given human impact in 27 interviews, these frequencies were used to rank human impacts into vulnerability scores. Numbers in parenthesis correspond to integer values only for aritmetic purposes of the model.

| **Human Activity** | **Stressors** | **N** | **Vulnerability score** |
| --- | --- | --- | --- |
| Agriculture | Non-point source organic pollution | 27 | High (4) |
| Aquaculture | Deforestation<br>Nutrients | 4 | Medium (3) |
| Centres of<br>population | Littering<br>Deforestation | 2 | Low (2) |

The weighted model of mangrove state (Figure 3) differs from the un-weighted model in revealing that mangroves closer to agriculture fields are more impacted than those adjacent to centres of population (pink coloured polygons in Figure 2B). The maximum impact score is reached in only one grid cell where there are aquaculture ponds in the south-east corner of the GoM.

**Figure 3.**
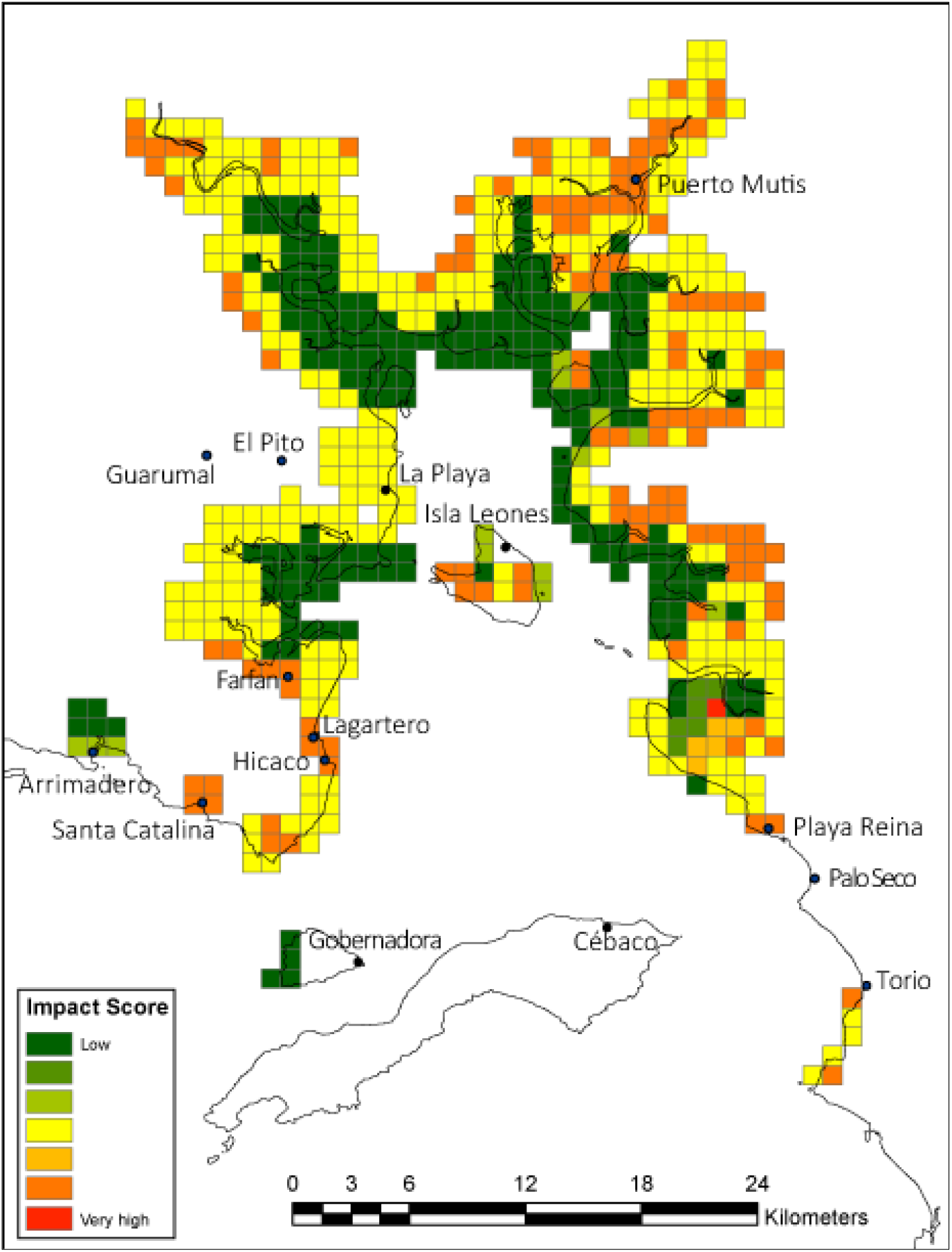
The Gulf of Montijo, Panamá. A cumulative impact model for human activities per 1km^2^ in the Gulf of Montijo showing levels of impact from low to high. This model has been weighted based on the vulnerability of mangroves to different human activities.

For the un-weighted and weighted impact scores, 30% of cells were classified as low impact, and 70% were medium to very high impact. Of the cells containing impacts, agriculture was the most common, present in 95% of cells, followed by centres of population present in 31% of cells, whilst aquaculture was present in only 2.7% of impact cells in the model. Supporting Information shows the detailed number of cells present in each impact category of the unweighted and weighted models.

## 4. Discussion

The GoM is a complex landscape of mangroves immersed in an active agricultural region with various human activities exerting influence on them. The impact score model showed that most of the mangroves of the GoM were affected by human activities (70%), with only 30% of cells containing mangroves without any directly adjacent impact. Cumulative impact maps of marine ecosystems have proven useful to inform managers on the implementation of ecosystem-based management, marine protected areas and ocean zoning (Halpern et al. 2008; Selkoe et al. 2009; Halpern et al. 2009). Ground-truthing allowed for evaluation of the ability of the impact score model to reliably assess the condition of mangroves in the GoM, in terms of the classification of human activities adjacent to mangroves. Google Earth has been shown to be a powerful, open-access tool for scientists and conservationists to assess the state of an ecosystem (Yu and Gong 2012). In this study, by using more than one source of land use information (i.e. satellite images in Google Earth, GIS layers, interviews, and ground-truthing), we avoided underestimation or misidentification of human impacts.

Our analysis shows that most of the mangroves of the GoM are influenced by human activities with agriculture being the most influential activity, present in 95% of the cells, followed by centres of population then aquaculture. The GoM is situated in the second largest agricultural region of Panamá where cattle farming, rice and sugar cane production contribute significantly to the local and national economy (ANAM 2004). However, despite restrictions on mangrove clearing, the area given to rice crops adjacent to mangroves has increased since 2000 (ANAM 2004). This information reflects recent findings which suggest that rice crops are responsible for the fastest rate of mangrove deforestation in Southeast Asia (Richards and Friess 2015). The damage to mangroves is not limited to direct deforestation. Agricultural crops grown close to mangroves are often ones that need a lot of water (e.g. melon, watermelon and rice), chemical pesticides, and fertilizers (Bach 2007). Agrochemicals and waste products discarded from farms, are transported by rivers and creeks, or are sluiced directly into mangroves and washed into estuaries, killing high numbers of organisms in the water and affecting fish catches (Trejos et al. 2007a; Kaufmann 2012).

In interviews, 100% of fishers said that agriculture posed the most important threat to mangroves and the fish populations they support. Fishers cited specific examples from their own experiences where agriculture has negatively affected mangroves in the GoM directly, because of clearance of mangroves for crops, and indirectly, because livestock husbandry and crop farming can result in toxic quantities of chemicals entering the sea. The scale of this agricultural runoff can be large in the GoM, because fertilization and application of pesticides are not only carried out with ground sprayers, but also from airplanes. Wetlands are known to be routinely contaminated by pesticides and fertilizers on adjacent agricultural areas (Alho and Vieira 1997; Donald et al. 1999; Hill 2003; Hernández-Romero et al. 2004), however the downwind drift from the aerial application has been estimated to be more than four times higher than that produced by ground sprayers (Ware et al. 1969). This might be because aerial fumigation has heightened effects on small streams and ponds, as they are hard to avoid when crops and wetlands share a common boundary (Hill 2003). Studies have shown that the aerial application of pesticides are directly lethal to wetlands wildlife, and that high mortalities are seen not only in fish given high levels of pesticide in water runoff (Hill 2003), but also in birds and mammals (Pimentel 2005). It has been shown that the concentration of pesticides in wetland waters is directly related to precipitation events (Donald et al. 1999), suggesting this may also be the case in the GoM, where large-scale fish mortality is usually observed at the start of the rainy season (Trejos et al. 2007b).

Some studies have shown changes in fish species assemblages following mangrove degradation or clearance (Williamson et al. 1994; Shervette et al. 2007; Shinnaka et al. 2007; Adite et al. 2013). When Adite et al. (2013) examined mangroves of varying degrees of degradation in West Africa, they found that fish species richness and diversity were significantly lower in degraded sites than in restored areas. In Thailand, Shinnaka et al. (2007) demonstrated that mangrove deforestation had marked effects on fish assemblages, as sites with mangroves had higher numbers of fish species and individuals than sites that had been cleared of mangrove forest.

In the GoM the majority of fishers recognized the importance of mangroves for their livelihoods, as seen elsewhere (MacKenzie 2001; Walters et al. 2008; Hussain and Badola 2010; UNEP 2014). Given the shortcomings of current fisheries data collection protocols in small-scale fisheries, the contributions of this sector to food security, local livelihoods, and poverty alleviation are likely to be undervalued in Panamá, as it is often the case in developing nations (Mills et al. 2011). One way to tackle this is the collection of fisheries data with household socio-economic data, as this provides a robust insight into the societal role of fisheries and ultimately contributes in a meaningful way to policy development (Mills et al. 2011).

The map of human impacts for the GoM showed that inland boundaries of mangroves were likely to be influenced by one or more human activities simultaneously, with cumulative consequences. Hence the protected area designation of the GoM, as a Wetland of International Importance, needs to adequately protect against indirect influences of peripheral land use activities. Our results highlight often overlooked impacts on mangroves from adjacent land use, and call for management strategies to regulate human impacts threatening mangroves resources, especially for the effects of fertilizers and pesticides on fishery species. For example, given that wetlands in agricultural landscapes are exposed to high levels of pesticides (Donald et al. 1999) fumigation should be regulated. This could be the first step towards a spatial planning approach that integrates management and legislation of multiple user groups and stakeholders (Crowder and Norse 2008) which is crucial to reduce the impacts that human activities cause on mangroves and fish stocks. This study provides a quantitative, spatially referenced assessment of the state of mangroves in the GoM within the context of the larger human landscape around them, highlighting the importance to regulate human activities and tailor management to the reality of the GoM.

## Acknowledgements

JLA thanks the following organizations for funding through fellowships: the Colombian Department of Science and Technology (COLCIENCIAS), the Schlumberger Foundation Faculty for the Future Programme, the World Wildlife Fund’s Russell E. Train Education for Nature program, the Smithsonian Tropical Research Institute (STRI), and Save our Seas Foundation (SOSF2013APP2N). Thanks to Dr A. Altieri, Dr J. Maté and M. Solano, from STRI for their helpful advice in different aspects of this research. Thanks to Fundación Marviva Panama for spatial data. Thanks to M. Villate and J.C. Cubillos from Fundación Talking Oceans for their assistance with interviews in the field

